# Direct Observation of the Mechanism of Antibiotic Resistance by Mix-and-Inject at the European XFEL

**DOI:** 10.1101/2020.11.24.396689

**Authors:** Suraj Pandey, George Calvey, Andrea M. Katz, Tek Narsingh Malla, Faisal H. M. Koua, Jose M. Martin-Garcia, Ishwor Poudyal, Jay-How Yang, Mohammad Vakili, Oleksandr Yefanov, Kara A. Zielinski, Saša Bajt, Salah Awel, Katerina Dörner, Matthias Frank, Luca Gelisio, Rebecca Jernigan, Henry Kirkwood, Marco Kloos, Jayanath Koliyadu, Valerio Mariani, Mitchell D. Miller, Grant Mills, Garrett Nelson, Jose L. Olmos, Alireza Sadri, Tokushi Sato, Alexandra Tolstikova, Weijun Xu, Abbas Ourmazd, John H. C. Spence, Peter Schwander, Anton Barty, Henry N. Chapman, Petra Fromme, Adrian P. Mancuso, George N. Phillips, Richard Bean, Lois Pollack, Marius Schmidt

## Abstract

In this study, we follow the diffusion and buildup of occupancy of the substrate ceftriaxone in *M. tuberculosis* β-lactamase BlaC microcrystals by structural analysis of the enzyme substrate complex at single millisecond time resolution. We also show the binding and the reaction of an inhibitor, sulbactam, on a slower millisecond time scale. We use the ‘mix-and-inject’ technique to initiate these reactions by diffusion, and determine the resulting structures by serial crystallography using ultrafast, intense X-ray pulses from the European XFEL (EuXFEL) arriving at MHz repetition rates. Here, we show how to use the EuXFEL pulse structure to dramatically increase the size of the data set and thereby the quality and time resolution of “molecular movies” which unravel ligand binding and enzymatically catalyzed reactions. This shows the great potential for the EuXFEL as a tool for biomedically relevant research, particularly, as shown here, for investigating bacterial antibiotic resistance.

**One Sentence Summary:** Direct observation of fast ligand binding in a biomedically relevant enzyme at near atomic resolution with MHz X-ray pulses at the European XFEL.

---

Combatting the rise of infectious diseases requires a collaborative and interdisciplinary approach. Structural biologists can contribute by investigating the reaction mechanisms of biomedically significant enzymes as a structural basis to develop cures for diseases. Bacterial infections with strains that are resistant to currently available antibiotics are on the rise (*1*). A study sponsored by the British government projected that in the near future more people will die from untreatable bacterial infections than from cancer (https://amr-review.org/). Bacterial enzymes that inactivate currently available drugs are central to antibiotic resistance (*2*) and unraveling the catalytic mechanism of these enzymes will be beneficial for the development of novel antibiotics (*3*). β-lactamases such as *M. tuberculosis* β-lactamase (BlaC), catalytically inactivate β-lactam antibiotics and are responsible for the emergence of multidrug and extensively drug resistant bacterial strains(*4*). Infectious diseases that could be treated with β-lactam antibiotics in the past may become untreatable.

With time-resolved crystallography, structures of intermediates and kinetic mechanisms can be extracted simultaneously from the same set of X-ray data (5, 6). Pioneering time-resolved crystallographic experiments at XFELs were all of the pump-probe type, where an optical laser pulse triggers a reaction in the crystallized molecules which are probed by x-ray pulses after a controlled delay (*7, 8*), with experiments capable of reaching sub-ps time resolutions (*9, 10*). Photoactivation, however, requires a light sensitive cofactor, a chromophore, located in the protein of interest to absorb the light. For investigating reactions in enzymes, this light absorption must trigger a reaction that either promotes catalysis directly (*11*) or adjusts the activity of the enzyme (*12–14*). Most enzymes, however, are neither activated nor regulated by light, meaning the technique can only be directly applied in a narrow range of cases. Broader application requires great effort and chemical expertise to either engineer photoactive enzymes or to design photoactive compounds that can by soaked into, and activated in, enzyme crystals (*15, 16*).

With the ‘mix-and-inject’ technique (*17–20*) photoactivation is not necessary. Substrate is rapidly mixed with small enzyme crystals during sample delivery (*21*). Mixing occurs at a well-controlled location ‘en route’ to the X-ray beam. During the time-delay ΔT_mi_ that occurs between mixing and injection, substrate diffuses into the crystals and binds to the enzyme. The complex formed by the substrate and the enzyme then initiates the enzymatic cycle. Variation of ΔT_mi_ allows measurement of chemical rate coefficients, and can associate an atomic-resolution structure to intermediate states which occur during protein reactions. This can reveal the mechanism of enzyme action at the molecular level, or the binding of a drug molecule. The feasibility of the mix-and-inject technique on enzymes was first demonstrated with the BlaC on longer millisecond timescales (*18, 19*). The observation of intermediate state structures, and maximization of the potential time resolution in both photoactivation and mix-and-inject techniques, relies on an accurately gauged start time of the reaction inside the crystals. In photoactivation experiments this requires a sufficient penetration of an optical laser into the crystal to ensure a reaction is simultaneously triggered in a significant fraction of the molecules. The laser power, however, must be carefully adjusted to avoid multi-photon excitation pathways and damage (*22, 23*). In mix-and-inject experiments, the diffusion time of substrate into the crystal limits the ability to discriminate diffusion and kinetics, including substrate binding. In both cases these considerations require micron or sub-micron crystal sizes, with limited scattering power.

At X-ray Free Electron Lasers (XFELs) small, micrometer (μm) and sub-μm sized, crystals can be examined due to the immense X-ray pulse intensity (*24*). Micro-crystals are destroyed by the pulses, and new crystals must be delivered to the X-ray interaction point in a serial fashion (*25*). Since the XFEL pulses are of femtosecond duration, diffraction patterns are collected before the crystals suffer significant radiation damage, resulting in X-ray structures that are essentially damage-free (*26, 27*) and suspended in their current reaction state. The combination of serial femtosecond crystallography (SFX) (*24, 25*) with mixing before injection has been denoted ‘Mix-and-Inject Serial Crystallography’ (MISC) (*18–20*).

The BlaC reaction with the cephalosporin antibiotic CEF is an excellent candidate for exploration with MISC. Previously, this reaction was investigated for ΔT_mi_ longer than 30 ms (*18, 19*). At 30 ms, however, the CEF binding sites in BlaC were essentially fully occupied (*19*), a state also reached on similar time-scales for other proteins and enzymes (*20, 28, 29*). The important substrate binding phase and the formation of the enzyme-substrate complex, however, remain elusive.

## Mix-and-Inject Experiments at the EuXFEL

To observe substrate binding directly, we performed a pioneering single ms MISC experiments at the SPB/SFX instrument(*30*) of the superconducting European XFEL (EuXFEL) in Germany(*31*). Here, we used the EuXFEL MHz pulse structure(*8*) to measure the binding of the large CEF substrate to BlaC on timescales much faster than 30 ms to capture diffusion and the binding kinetics. In addition, we investigated the reaction of the BlaC with an inhibitor, sulbactam (SUB) on a millisecond time scale. The biochemistry of SUB and its application in combination with β-lactam antibiotics are described in detail elsewhere (*32*). SUB binds to the active site of BlaC and reacts with the catalytically active serine of β-lactamases to form various covalently bound species. Most abundant is the so-called *trans*-enamine (*trans*-EN) species (Scheme S1, compound III) that inhibits β-lactamases and helps to eliminate β-lactamase induced antibiotic resistance. The static structures of *trans*-ENs with β-lactamases, including BlaC, were recently characterized (*33, 34*) but structures of the early species that form during SUB binding remain elusive.

BlaC crystals grown with ammonium phosphate form platelet shaped microcrystals (Fig. S1 a) which are ideal for mix-and-inject investigations on fast time scales. Fig. 1 a shows the structure of the BlaC in this crystal form. A dense suspension of microcrystals was injected into the ~3 μm X-ray focus of the SPB/SFX instrument (*30*) of the EuXFEL (Fig. 2). For CEF, three time delays ΔT_mi_ at 5 ms, 10 ms, and 50 ms were probed. The reaction of BlaC with SUB was probed 66 ms after mixing. As a reference and a control the BlaC crystals were mixed with water and probed 10 ms after mixing (see Table S1 for data statistics and Material and Methods for further details).

**Figure 1.**
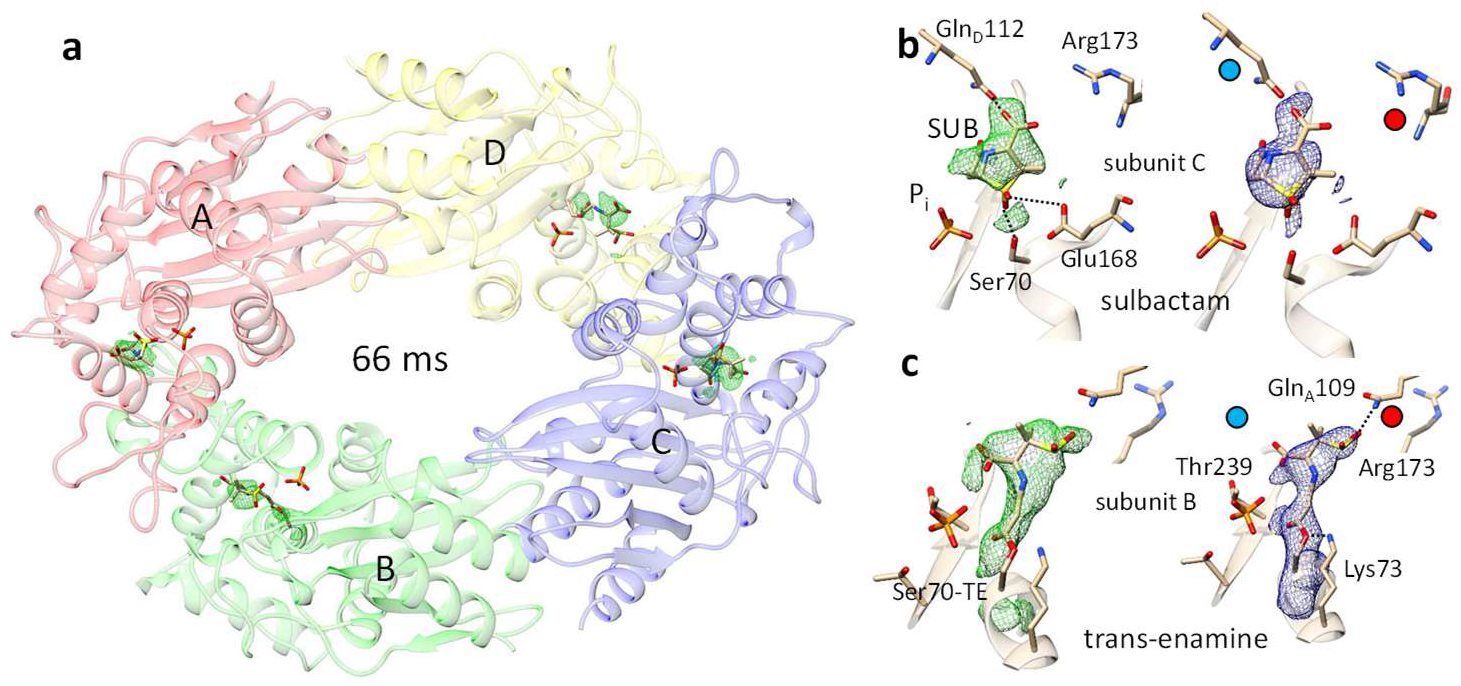
The crystal structure of BlaC. (a) Sulbactam (SUB) binds to all four subunit (A-D) in the asymmetric unit. The phosphates in the active sites are not replaced. Isomorphous difference map (DED_iso_) (2 sigma) shown in green. (b) Active site in subunit C with non-covalently bound intact sulbactam, left side: DED_iso_ map, right side: 2mFo-DFc map after refinement. Close by amino acids and the phosphate (P_i_) are marked. (c) Active site in subunit B with *trans*-enamine bound to Ser-70, left side: DED_iso_ map, right side: 2mFo-DFc map after refinement. Red and blue dot show important differences between the subunits. Gln112 from the adjacent subunit is not located close-by, and the Arg173 is extended in subunit B leaving subunit B more accessible to ligands and substrate.

**Figure 2.**
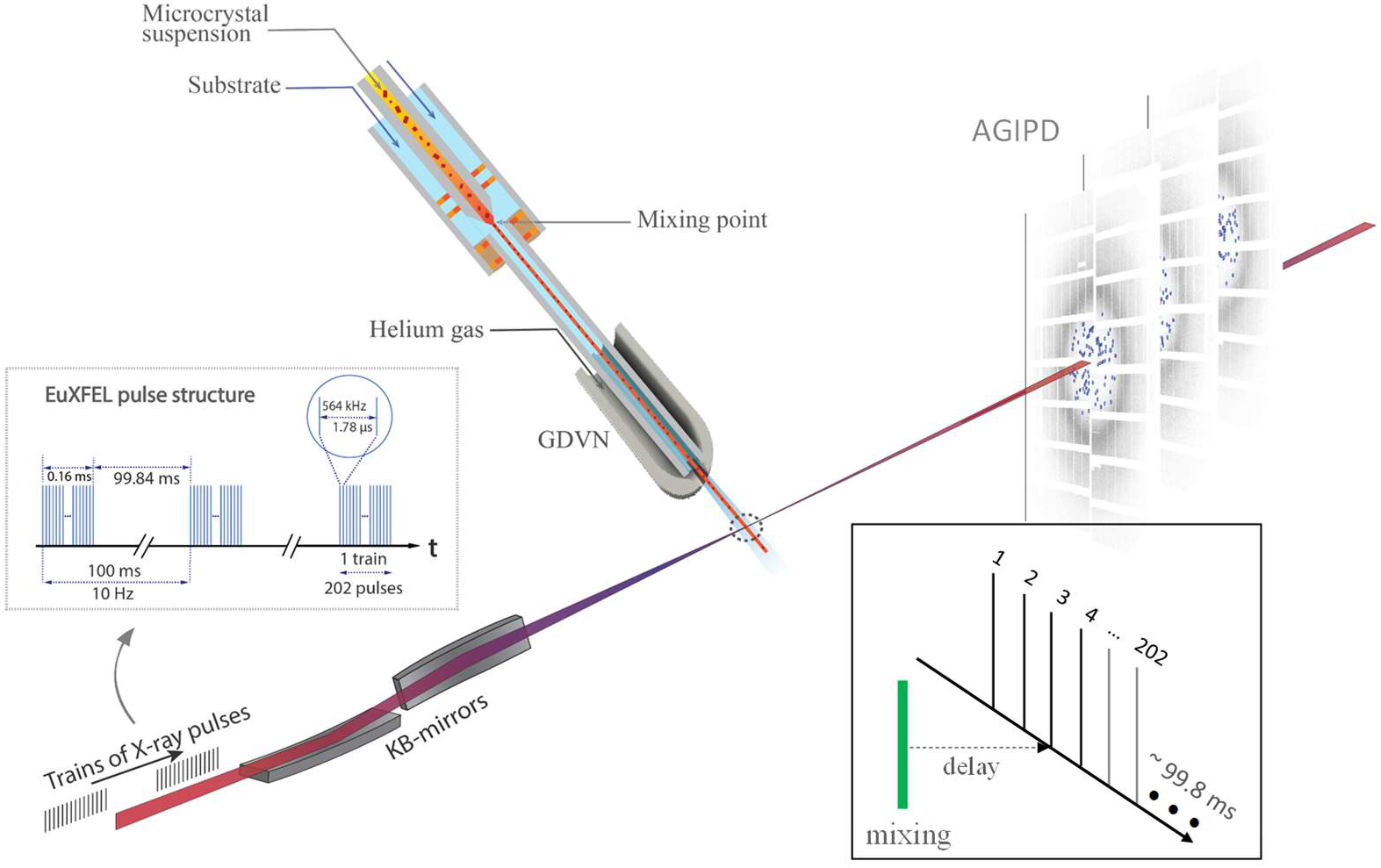
Experimental setup at the European XFEL. Microcrystals are mixed with substrate, and injected into the X-ray after a delay determined by the distance between the mixing region and the X-rays, the capillary width, and the flow rate. Diffusion of substrate into the crystals occurs during this time. The mixture is probed by trains of X-ray pulses. The trains repeat 10 times per second. Pulses within the trains repeat with 564 kHz, hence the pulses are spaced by 1.78 μs. 202 pulses were in each train for this experiment. The AGIPD collects the diffraction patterns and reads them out for further analysis. Insert, data collection: With a once selected injector geometry and flow rate, the delay is fixed by the distance of the mixing region from the X-ray interaction region. All pulses in all trains (here pulse #3) probe the same time delay. The EuXFEL pulse structure is most efficiently used.

## The Induced Fit: Formation of the Enzyme Substrate Complex

Fig. 3 shows CEF binding. As observed in a previous study at the longer ΔT_mi_ of 2 s (*18*), CEF binds only to subunits B and D. In Fig. 3 a, DED_omit_ in the active site of subunit B is shown. On early time scales (5 ms and 10 ms after mixing) we observe a crystal-averaged electron density of a CEF and a phosphate (P_i_) molecule at this position. The P_i_ is also found near the CEF binding site in the unliganded (unmixed) form. At a ΔT_mi_ of 50 ms, the P_i_ density has vanished. Occupancies for P_i_ and CEF were refined using the program ‘Phenix’(*35*) at all ΔT_mi_ (Table S2). At the 5 ms delay, the P_i_ and the CEF occupancies are both approximately 50 %. The available catalytic sites in subunits B and D are equally occupied either by a CEF or by a P_i_ (Fig. 4 a). At ΔT_mi_ = 50 ms the P_i_ occupancy refines to 19 %, that of the CEF to 82 %. In agreement with previous work(*19*), an additional CEF molecule is identified close to each active site that weakly interacts (stacks) with the already bound CEF bound there (Fig. S2). The stacking sites are only transiently visited by CEF molecules until the active sites are fully occupied. Unit cell parameter changes roughly follow CEF binding and P_i_ release (Table S1, Fig. 4 a, insert). When the about 2.5 times smaller sulbactam binds, the P_i_ is not replaced, and the unit cell parameters do not change (Table S1). We postulate that the size of the ligand, as well as the displacement of the strongly negatively charged P_i_ may contribute to the unit cell changes as observed when CEF is mixed in.

**Figure 3.**
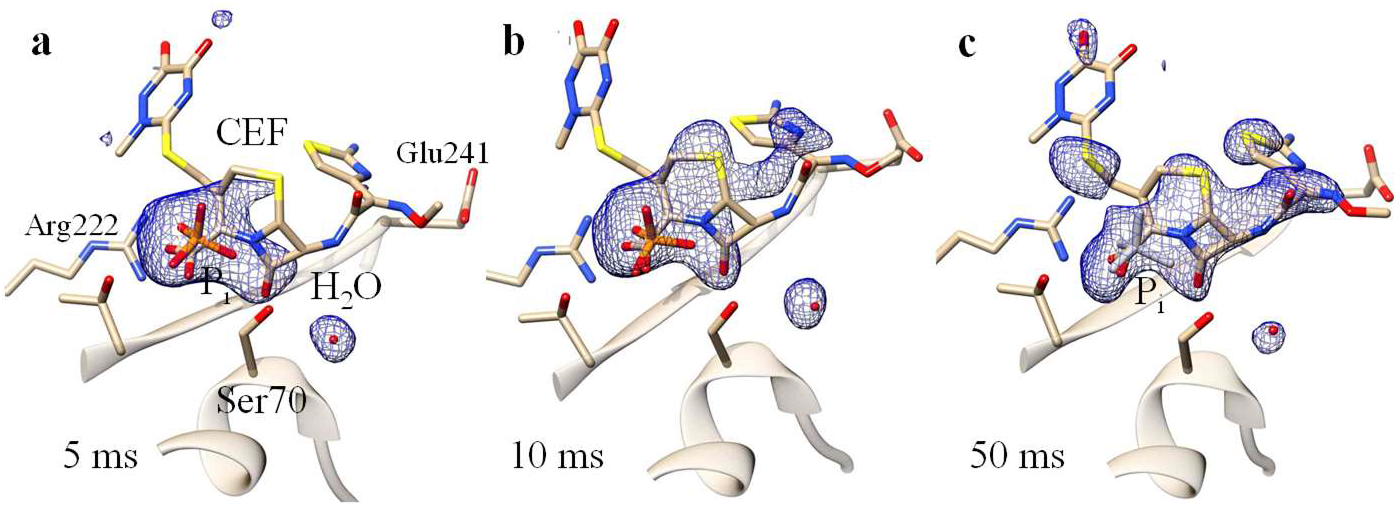
Omit difference electron density in the active center of BlaC, subunit B. (a) 5ms, (b) 10 ms after mixing and (c) 50 ms after mixing. Ser-70, CEF, the phosphate (P_i_) and the water molecule are marked. The P_i_ is essentially absent in (c). Some nearby amino acids are displayed in addition.

**Figure 4.**
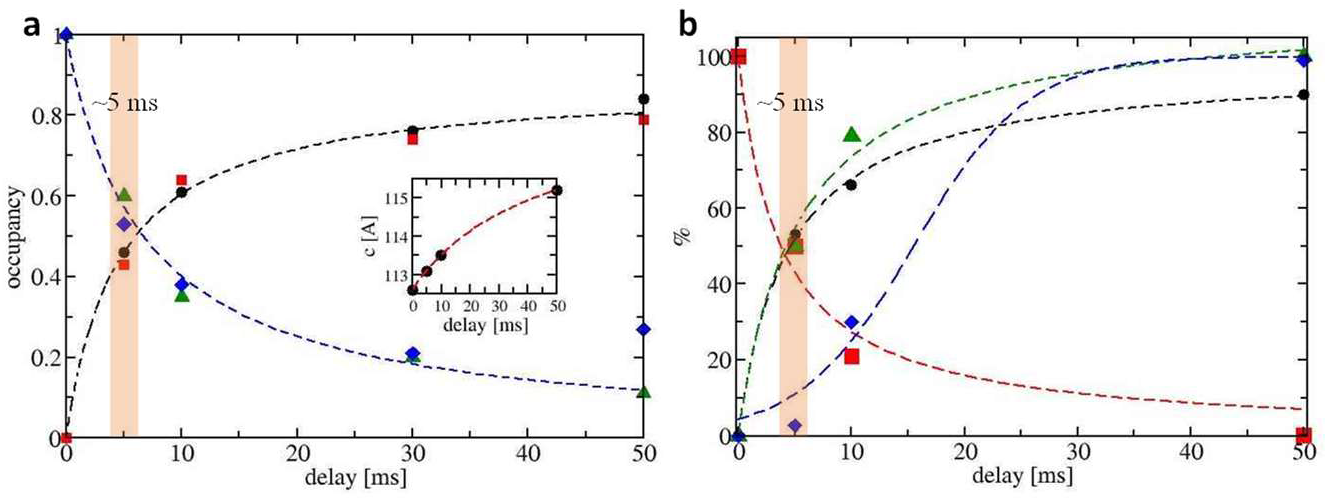
BlaC, CEF and the BlaC-CEF complex as a function of time. (a) Occupancies of CEF at 5 ms, 10 ms and 50 ms in subunits B (spheres) and D (squares) as well as those of phosphate (P_i_) (green triangles and blue diamonds) are plotted as a function of delay. The data are fitted with saturation curves (eqs. S1 and S2, black and blue dashed lines). The two curves intersect at around 6 ms. Insert: corresponding change of unit cell axis c. (b) Concentrations (in %) as calculated from diffusion and binding (eqs. S1 - S5). Black spheres: increase of the free BlaC concentration, red squares: decrease of the free BlaC concentration, green triangles: increase of the BlaC-CEF complex concentration averaged over all voxels in the crystal, blue diamonds: increase of the BlaC-CEF complex in the center of the platelet shaped crystals. Data were fitted by saturation curves (corresponding black, red and green dashed lines). Concentrations in the crystal center (blue dashed line) were fitted by a logistic function.

As more CEF binds, Ser-70 moves towards the P_i_ position, and the P_i_ is replaced at the same time (Table S3 c). Other amino acids such as Asn 172 and Asp 241 move closer to the CEF. We can now develop a mini-movie for the formation of an enzyme-substrate (ES) complex. This movie shows an induced fit (Supplementary Movie 1) and visualizes in real time how the active site closes around the CEF. The induced fit phase finishes after ΔT_mi_ = 10 ms when the CEF occupancy approaches saturation. In the inactive subunits A and C, two glutamines, Gln-109 and Gln-112, from adjacent, non-crystallographically related subunits, extend into the active sites and would clash severely with the dioxo-triazine ring on one side and with the amino-thiazole ring on the other side of CEF (Scheme S1, compound I), thus effectively preventing CEF binding.

The ES complex formation is most important since it triggers the enzymatic cycle. Hence, it determines the time resolution of the MISC method. The ES complex consists of CEF non-covalently bound in the active site of BlaC (Scheme S2). CEF is delivered by diffusion into the crystals. Crystals must be small enough to enable short enough diffusion times, so that the binding kinetics can be observed. However, MISC does not measure the free substrate concentration in the crystals, and therefore diffusion is rather observed indirectly through the increase of occupancies of well-ordered substrate molecules in the active centers of BlaC. When the diffusion times are very short, occupancies may accumulate on a timescale longer than the diffusion time, as they are governed by the binding kinetics. The ES complex formation is therefore not only dependent on the ligand concentration delivered by diffusion but also on the magnitude of the rate coefficients in the kinetic mechanism.

On the short timescales employed here, the formation of later intermediates and the catalytic turnover of the BlaC do not play a role. Both processes unfold over much longer timescales than the time delays examined here (*19*). Diffusion follows Fick’s 2^nd^ law from whose solution the CEF concentration can be estimated at any position in the crystal and at any time. CEF binding to the active sites of BlaC is dependent on the free BlaC concentration in the crystal, the CEF concentration and the rate coefficients that describe the kinetic mechanism. There is only one free parameter, the diffusion coefficient D that can be inferred by matching calculated occupancies to the occupancies measured at the ΔT_mi_ (compare Fig. 4 a and Fig. 4 b, see Methods for details). CEF diffusion is about a factor of twelve slower (D_eff_ ~ 2 × 10^−7^ cm^2^/s) within the BlaC crystals compared to water. This slowdown is in agreement with findings that were previously obtained from simulations on substrate diffusion in enzyme crystals (*36*). Fig. 5 depicts the result of the calculation. Estimates of enzyme-ligand occupancies are now directly deduced from time-resolved X-ray crystallography everywhere in a crystal after mixing. Not surprisingly, at 5 ms the occupancy is high (> 90 %) only near the crystal surface where enough substrate is present to promote ES formation with a high rate. In the center of the crystals the ES complex concentration is initially small (Fig. 4 b, Fig. 5 a). The binding rate is not sufficiently high to generate significant occupancy. After ΔT_mi_ = 10 ms the binding rate increases, until at 30 ms full occupancy of the BlaC-CEF complex is reached (Table S5 b and Fig. 4 b, green dashed line) everywhere. With the rapid diffusion of CEF into small BlaC crystals we are now able to quantify variations of substrate, enzyme and ES concentrations across the enzyme crystal volume (Table 1, Fig. 4 and Fig. 5). The remarkable speed of the ES accumulation shows that the mix-and-inject technique can be used to characterize enzymes with turnover times much faster than that of BlaC. The direct observation of the important initial ligand and substrate binding phase in biomedically relevant enzymes is possible.

**Figure 5.**
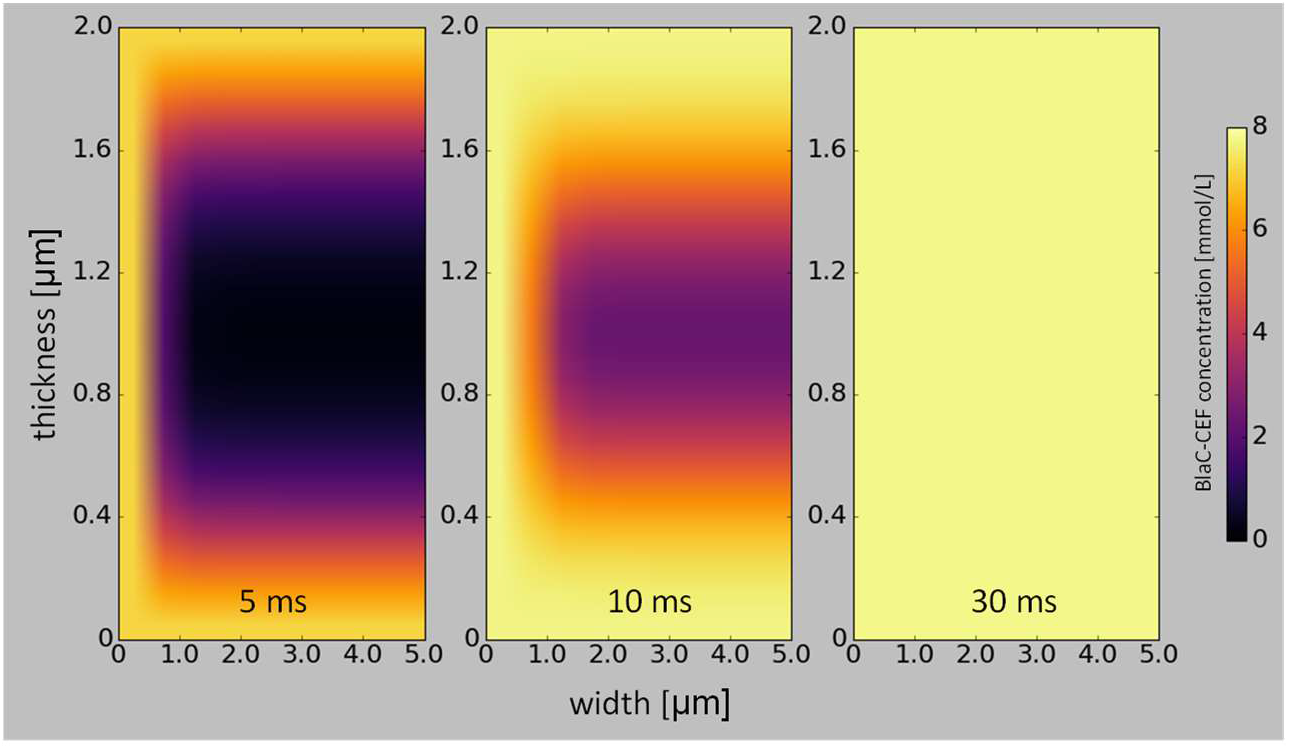
Concentrations of the BlaC-CEF complex in 10 × 10 × 2 μm^3^ platelet shaped crystals, (a) 5 ms, (b) 10 ms, and (c) 30 ms after mixing with 200 mmol/L ceftriaxone (150 mmol/L final concentration assumed). The concentrations are shown in different colors (see scale bar on the right) in central cross sections through half the width of the crystals. The drawings are not to scale, since the sections displayed are 5 μm horizontally (width) and 2 μm vertically (thickness). The enlargement along the short 2 μm axis allows the display of nuanced occupancy differences.

**Table 1.**
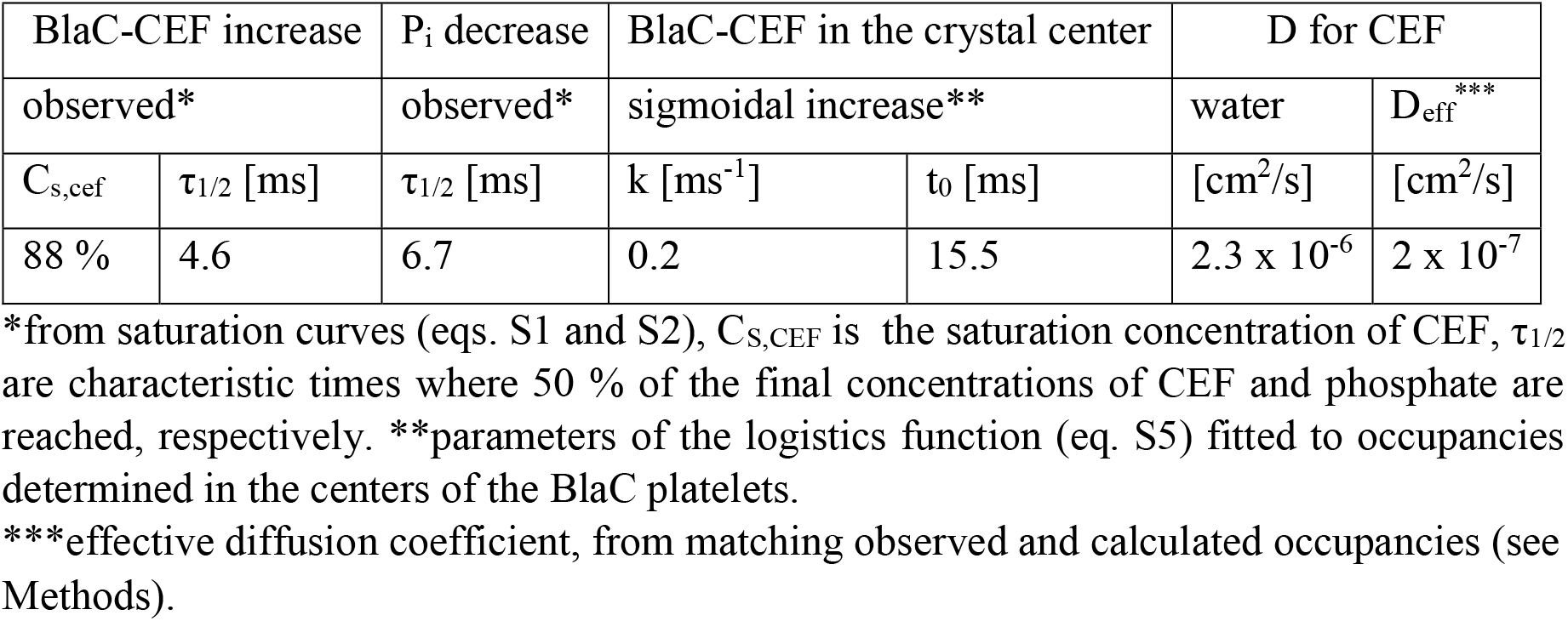
BlaC-CEF complex formation in microcrystals. Results from comparing observed and calculated occupancies.

| BlaC-CEF increase |  | P<sub>i</sub> decrease | BlaC-CEF in the crystal center |  | D for CEF |  |
| --- | --- | --- | --- | --- | --- | --- |
| observed* |  | observed* | sigmoidal increase** |  | water | D<sub>eff</sub>*** |
| C<sub>s,cef</sub> | τ<sub>1/2</sub> [ms] | τ<sub>1/2</sub> [ms] | k [ms<sup>-1</sup>] | t<sub>0</sub> [ms] | [cm<sup>2</sup>/s] | [cm<sup>2</sup>/s] |
| 88 % | 4.6 | 6.7 | 0.2 | 15.5 | 2.3 x 10<sup>-6</sup> | 2 x 10<sup>-7</sup> |

\*from saturation curves (eqs. S1 and S2), C<sub>s,CEF</sub> is the saturation concentration of CEF, τ<sub>1/2</sub> are characteristic times where 50 % of the final concentrations of CEF and phosphate are reached, respectively. \*\*parameters of the logistics function (eq. S5) fitted to occupancies determined in the centers of the BlaC platelets.
\*\*\*effective diffusion coefficient, from matching observed and calculated occupancies (see Methods).

Since the ES complex (here the BlaC-CEF complex) triggers the enzymatic cycle, accurate kinetics can be extracted to the point that the time required to accumulate sufficient ES-complex approaches the life-time of the next intermediate in the catalytic cycle (*17*). This finding holds for any other technique (*15*) which aims to trigger enzymatic reactions, even in non-crystalline samples. Not only it is required to bring enough substrate near the vicinity of the enzyme, but also the binding kinetics needs to be taken into account. With microcrystals below a certain crystal size, binding of substrate, and not the diffusion of substrate into the crystal volume, may become rate limiting. As a consequence, for BlaC crystals about a size of 1 μm, the speed of the ES complex formation is not substantially different from that in solution. The Deff determined here suggests that accurate measurements of the substrate binding kinetics would not be possible with significantly larger crystals. Enzymes with turnover times faster than the BlaC will usually also display faster substrate binding kinetics with larger k_on_ rate coefficients. In such cases, the crystal sizes (and their size distributions) or perhaps temperature must be adjusted appropriately to ensure that the diffusion times can catch up with the substrate binding rates.

## Inhibitor Binding and Reaction

SUB binds to all four subunits in the asymmetric unit, which is in stark contrast to CEF that binds only to subunits B and D. Notably, however, the DED_iso_ is different for subunits A, C and B, D. In subunits B and D, the electron density can be interpreted with a sulbactam *trans*-EN covalently bound to Ser-70 (Scheme S1, compound III, Fig. 1 c, Table S2). The orientation of the *trans*-EN in the active site is determined by residues Lys-73 and Thr-239 and in particular by Gln-109 from the adjacent non-crystallograpically related subunit (Table S3 b). In subunits A and C, the DED_iso_ shows changes in a region more distant from Ser-70. Electron density details extend from a spheroidal, central DED feature. An intact SUB molecule that is non-covalently bound to the active site (Fig. 1 b, Scheme S1, compound II) can be fitted. The SUB is oriented so that the ring sulfur-dioxide points towards Ser-70 with the β-lactam ring pointing away. We hypothesize that this ‘up-side-down’ orientation is enforced by Arg-173 and Gln-112 (Table S3 a), where Gln-112 is protruding deep into the active sites from the adjacent, non-crystallographically related subunits (Fig. 1 b).

The diffusion time is fast enough that 66 ms after mixing all non-covalently bound SUB molecules in subunits B and D have reacted to the covalently bound *trans*-EN. This is quite unexpected as it was suggested that it would take minutes for the enamine to form after binding of SUB in the BlaC (*32–34*). Although the SUB binds non-covalently to the active sites in subunits A and C, the reaction with the Ser-70 did not occur within ΔT_mi_ = 66 ms. It may be that SUB will not react further in these subunits. It could also be that the structure of the BlaC-SUB complex is an interesting intermediate on the reaction pathway to the *trans*-EN. Experiments exploring longer ΔT_mi_ will clarify this situation. In subunits B and D Glu-112 is not near the active site, and Arg-173 displays a stretched, open conformation allowing the SUB to orient correctly towards the Ser-70, and to react further to the *trans*-EN that then non-competitively inhibits the BlaC (*34*). The near-by P_i_, which is displaced when the much larger ligand CEF is present, stays in place in all subunits and likely adds to the stability of both complexes.

## Protein Dynamics at the EuXFEL

In order to further investigate CEF and SUB binding and their reactions with Ser-70, a time series should be collected that consists of datasets at multiple ΔT_mi_ that span from a few ms to seconds. To achieve this, the EuXFEL pulse structure must be exploited most efficiently. Every X-ray pulse in all pulse trains provides observations of the same time delay, and our experiments took maximum advantage from the high pulse rate (Fig. 2, insert). This is in contrast to optical pump-probe experiments that require appropriate waiting times between the laser excitations to guarantee that the laser excited volume exits the X-ray interaction region, so that multiple laser activations can be avoided (*8*). We showed that diffraction data sufficient for good quality structure determination can be collected in about half an hour as demonstrated for the 50 ms CEF time point (Table S4). This time can be reduced substantially by limiting the number of diffraction patterns per dataset and by optimizing the crystal density flown through the mixing device. High crystal density will lead to higher hit rates but might also causes frequent interruptions caused by injector clogging. For our experiments, a fine balance between crystal size and crystal density was found so that the mix-and-inject experiments with CEF and SUB could be completed successfully with acceptable hit rates given the high X-ray pulse repetition rate at the EuXFEL. Previous experiments have shown that the collection of sufficient patterns for structure determination should be possible in less than 20 minutes at the detector-limited repetition rate of the EuXFEL (*8, 37*). This provides the tantalizing possibility to directly characterize the kinetic processes in biomolecules from single digit millisecond to longer time scales, within relatively short experimental times. The kinetics can rapidly change when environmental conditions are varied. It may be possible, for example, to control the temperature in the mixing injector delay line to determine barriers of activation from the resulting X-ray data (*38*). The full analysis of such a multi-dimensional data set requires the development and deployment of user-friendly classification algorithms to separate mixtures into their pure components (*39*) and derive kinetics and energetics (*38*) consistent with the electron density maps and structures of intermediate states along the reaction pathway.

Our experiments permitted a real-time view into the active sites of an enzyme during substrate and inhibitor binding. They facilitate more mix-and-inject experiments at the EuXFEL with unprecedented data collection rates allowing for more structures to be determined per allocated experimental time. This capability will become an important tool for biomedically relevant research in the years to come.

## Acknowledgements

We are very grateful to those who supported this experiment by being present in person at the European XFEL during the onset of the COVID-19 pandemic in March 2020. We acknowledge the European XFEL in Schenefeld, Germany, for provision of X-ray free-electron laser beamtime at scientific instrument SPB/SFX and would like to thank the staff for their assistance. This work was supported by the National Science Foundation Science and Technology Center “BioXFEL” through award STC-1231306, and in part by the US Department of Energy, Office of Science, Basic Energy Sciences, under contract DE-SC0002164 (A.O., algorithm design and development) and by the National Science Foundation under contract numbers 1551489 (A.O., underlying analytical models) and DBI-2029533 (A.O., functional conformations). This material is based upon work supported by the National Science Foundation Graduate Research Fellowship Program to J.L.O. under Grant No. 1450681. The work was also supported by funds from the National Institutes of Health grants R01 GM117342-0404. Funding and support are also acknowledged from the National Institutes of Health grant R01 GM095583, from the Biodesign Center for Applied Structural Discovery at ASU, from the National Science Foundation award No. 1565180, and the U.S. Department of Energy through Lawrence Livermore National Laboratory under contract DE-AC52-07NA27344. K. A. Z. was supported by the Cornell Molecular Biophysics Training Program (NIH T32-GM008267). This work was also supported by the Cluster of Excellence “CUI: Advanced Imaging of Matter” of the Deutsche Forschungsgemeinschaft (DFG) -EXC 2056-project ID 390715994. CFEL is supported by the Gottfried Wilhelm Leibniz Program of the DFG; the project “X-probe” funded by the European Union’s 2020 Research and Innovation Program under the Marie Sklodowska-Curie grant agreement 637295; the European Research Council, “Frontiers in Attosecond X-ray Science: Imaging and Spectroscopy (AXSIS)”, ERC-2013-SyG 609920; and the Human Frontiers Science Program grant RGP0010 2017. This work is also supported by the AXSIS project funded by the European Research Council under the European Union Seventh Framework Program (FP/2007-2013)/ERC Grant Agreement no. 609920. The structures and diffraction data of the BlaC, unmixed and mixed with Ceftriaxone at 0 s, 5 ms, 10 ms and 50 ms are deposited to the Protein Data Bank (pdb) with the following access codes. CEF: 7K8L (unmixed), 7K8E (5 ms), 7K8F (10 ms), 7K8H (50 ms), SUB: 7K8K (66 ms). The SFX stream files were deposited in the CXIDB with code 170.

## Author Contributions

S.P., T.M., J.M-G., J.H.-Y., F.K., I.P., M.M., R.J., M.S., W.X., J.O. expressed, purified and crystallized the protein. R.B., K.D., H.K., M.K., J.K., G.M., T.S., M.V. operated the SPB/SFX instrument. L.P., A.M.K, G.C., K.A.Z. designed and provided injector nozzles. M.V., F. K., M.K. assembled and operated the nozzles, F.K., J.M-G., J-H.Y., L.G., P.S., collected the data. S.P., I.P., O.Y., V.M., P.S., A.T., A.B. processed the data. S.P., I.P., T.M., G.P. M.S. analyzed the data. I.P., M.F., A.S., F.K., P.S., M.S. logged the experiment. R.B., A.P.M., M.S.,G.P. designed the experiment. S.P., P.F., A.P.M., R.B., G.P., M.S. wrote the manuscript with input from all other authors.

## Supplementary Materials

**(The Supplementary Material is available from the authors on request)**

Materials and Methods

References 40 - 49

Schemes S1 to S2

Figs. S1 to S2

Tables S1 to S5

**Other Supplementary Material for this manuscript includes the following:** movie_1

